# MetaPathways v3.5: Modularity and Scalability Improvements for Pathway Inference from Environmental Genomes

**DOI:** 10.1101/2024.06.04.597460

**Authors:** Ryan J. McLaughlin, Tony X. Liu, Tomer Altman, Aditi N. Nallan, Aria S. Hahn, Julia Anstett, Connor Morgan-Lang, Kishori M. Konwar, Steven J. Hallam

## Abstract

Over the past decade MetaPathways has advanced as a modular pipeline for constructing environmental pathway genome databases (ePGDBs), increasing our understanding of microbial metabolism at the individual, population and community levels of biological organization. With this release, we have addressed several user experience issues related to installation, module integration, and database management. With a refactored code base, MetaPathways v3.5 enhances the user experience through streamlined installation via package indexes or containers, refined modules, and interface upgrades. It boasts updated algorithm support for sequence feature prediction, annotation, metabolic inference, and coverage metrics including genome resolved metagenomes. Tested and refined on synthetic datasets, MetaPathways v3.5 demonstrates improved performance and usability; facilitating more in-depth exploration of microbial interactions and metabolic functions in environmental genomes that scales with contemporary sequencing throughput.

**Availability and Implementation:** MetaPathways v3.5 is available *via* Anaconda, Docker, and Apptainer. The source code is available on BitBucket:https://bitbucket.org/BCB2/metapathways/ The documentation is available via ReadTheDocs:https://metapathways.readthedocs.io

## 1 Introduction

MetaPathways ([18]) is a modular pipeline originally designed to construct environmental Pathway/Genome Databases (ePGDBs) compatible with the editing and navigation features of Pathway Tools ([12]). MetaPathways’s capacity to chart microbial metabolism at different levels of biological organization supports insightful studies of microbial interactions in natural and engineered environments ([19, 28, 7, 22, 24, 5]) and has led to the development of community resources including the Encyclopedia of Environmental Genomes (https://engcyc.org) and GutCyc ([4]).

Since its inception, MetaPathways has been accumulating new features and algorithmic improvements, transforming a purpose-built pipeline into an increasingly advanced system for large-scale environmental genomic data processing and analytics ([6, 17, 23]). MetaPathways ’sarchitecture, although designed for continuous improvement, has faced several challenges due to increasing interdependence among modules and the growing complexity of managing the installation of third-party software and large reference databases. To address these issues, we present MetaPathways v3.5: the latest pipeline iteration introducing automated installation methods *via* package indexes or containers, refactored and disentangled modules, internal reference database management, and intuitive command line interface (CLI) upgrades, thus mitigating key barriers to entry and successful utilization.

## 2 Methods and Implementation

### 2.1 Sequence Feature Prediction

At the core of MetaPathways is a set of operations needed to predict gene and pathway features on quality-controlled assemblies. Feature prediction includes identification of open reading frames (ORFs), including ribosomal sub-units (rRNAs), and transfer RNAs (tRNAs) *via* Prodigal v2.6.3 ([8]), BARRNAP v0.9 ([29]), and tRNAscan-SE v2.0.12 ([1]), respectively. To further improve the performance of this module, an available wrapper tool for Prodigal, called pProdigal v1.0.1 was employed which allows for multi-threading for ORF prediction ([9]). Additionally, we built a custom wrapper to perform a similar function for tRNAscan-SE, called ptRNAscan, to provide fully parallel execution. We also added support to allow for more easy identification of overlapping ORFs, including rRNAs, and tRNAs.

### 2.2 Functional and Taxonomic Annotation

Predicted sequence features are annotated using one of two sequence alignment programs: NCBI BLAST ([31]) or FAST ([16]). FAST is a multi-threaded, input/output-optimized seed-and-extend alignment algorithm for nucleotide and protein sequences. By default, FAST is used for functional annotations and BLAST for taxonomic annotations. We added support for optional serial searches of multi-volume databases (to avoid loading all volumes into memory simultaneously), enabling MetaPathways to run in memory-constrained contexts, such as grid environments, individual workstations, and laptops. Lastly, we’ve updated support for calculating coverage statistics. This includes the use of the CoverM v0.6.1 software suite ([30]) to calculate various contig-level statistics such as raw count, RPKM, and TPM. Binary alignment map files (BAMs) ([20]) from CoverM are then used with custom code and the featureCounts v2.0.6 tool ([21]) to provide the same statistics for predicted features (*i*.*e*., ORFs, including tRNAs, and rRNAs).

### 2.3 Population and Community-level Pathway Inference

Pathway Tools is a software system supporting construction, management and navigation of Pathway/Genome databases (PGDBs) [13, 14, 12, 15], generated using the PathoLogic algorithm ([11]). A PGDB encodes knowledge related to genomic potential, metabolic network properties, and regulatory processes within cellular organisms. MetaCyc is a highly curated, non-redundant, and experimentally validated database of metabolic pathways representing all domains of life within the BioCyc Collection of PGDBs used as a trusted source for ORF annotation and metabolic pathway inference ([2, 11, 23]). The latest iteration of MetaPathways introduces redesigned utilities improving pipeline performance and the user experience. This includes utilities that enhance the extraction of Enzyme Commission (EC) numbers and MetaCyc reaction identifiers from supported reference databases, and the execution of the PathoLogic module from Pathway Tools on MetaPathways outputs. It now supports the creation of both population- and community-level ePGDBs by integrating the software application MAGsplitter ([25]), which accepts MetaPathways functional annotation tables along with mapping files for assigning contigs to metagenome assembled genomes (MAGs). MAGsplitter produces inputs for Patho-Logic to allow for population-level ePGDB to be built for each MAG. Additionally, we have updated and created new output tables, such as those mapping pathways to ORFs, thereby making it easier for users to correlate genomic sequence information with metabolic functions and coverage statistics.

**Figure 1:**
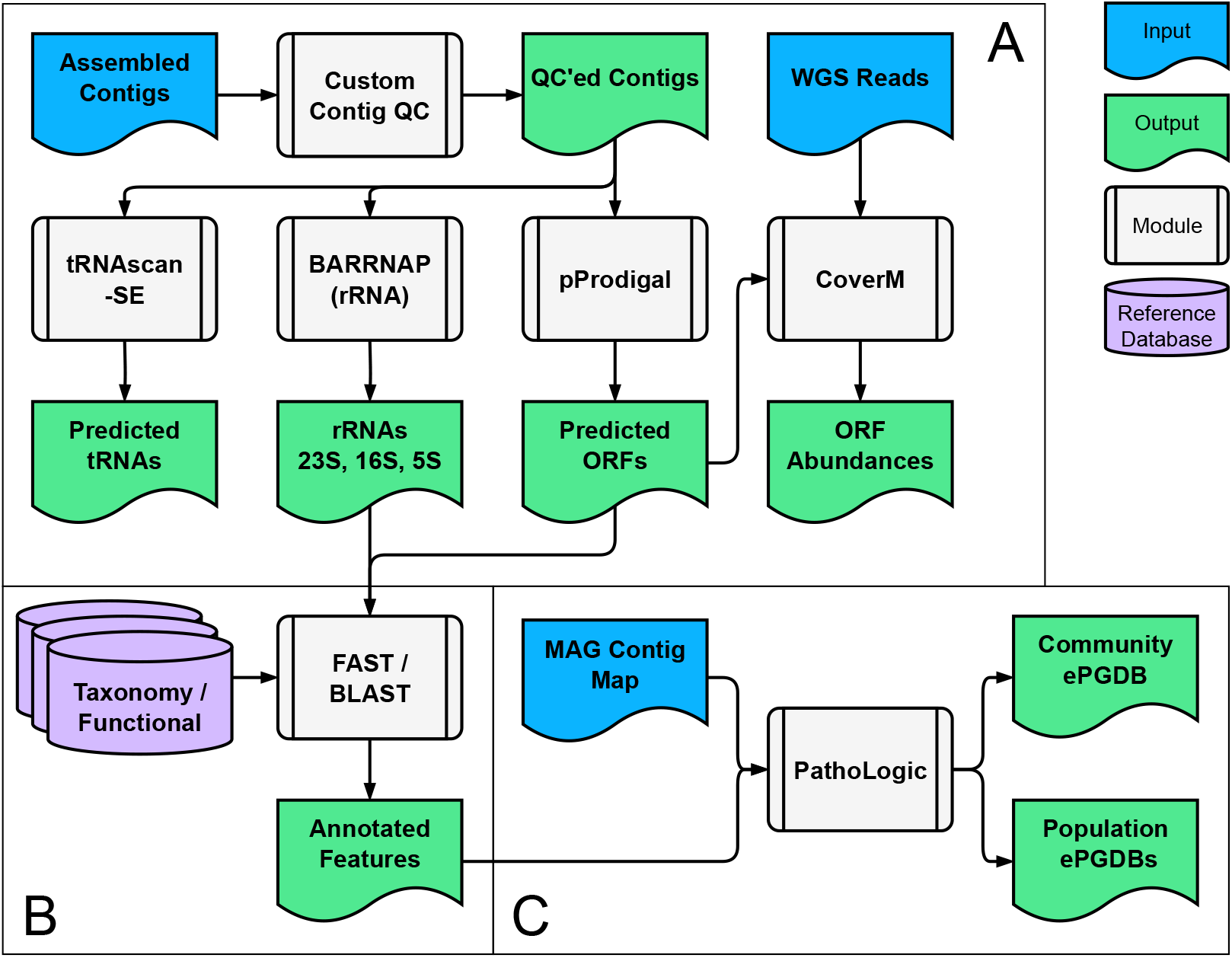
Modules of the MetaPathways workflow. **A:** Sequence feature prediction. **B:** Functional and taxonomic annotation. **C:** Community and population-level pathway inference.

## 3 Results

### 3.1 Performance

To provide performance metrics, we annotated the Critical Assessment of Metagenome Interpretation (CAMI2) Human Microbiome challenge synthetic communities ([26]) using default parameters and the SwissProt (release 2023 05) and MetaCyc (version 27.1) reference databases. Additionally, we produced MAGs using MetaBAT2 v2.15 ([10]) to allow for usage metrics for population-level pathway inference. We employed Pathway Tools (v27.0) to infer pathways for all metagenomes and MAGs. All analyses were performed using a virtual machine (VM) on the Genome Canada Arbutus Cloud ([3]), utilizing Intel Xeon Processors (Skylake, IBRS) with 16 cores, 176 GB memory, and running the Ubuntu 20.04.6 LTS GNU/Linux operating system. Each job used 16 cores and were run in series. Note that Pathway Tools does not make use of multi-threading and therefore, number of threads does not pertain to that part of the analysis.

A total of 15 metagenomes and 622 MAGs across five body sites were run to completion. Average runtime for the FAST/BLAST module was 168.19 ± 95.97 minutes (mins) with average memory usage of 12.53 ± 0.13 gigabytes (GBs). Maximum and minimum resource usage were 374.80 mins / 12.78 GBs and 52.80 mins / 12.34 GBs, respectively. This equates to a processing rate of approximately 1.6 megabases (MB) per minute and relatively stable memory usage across a wide range of assembly sizes, averaging 265±157 MB. After annotation, MAGs were mapped onto cognate assemblies to prepare Pathway Tools input files for ePGDB construction. Average resource usage for this step was 1.73 ± 1.09 mins / 1.17 ± 0.88 GBs. Pathway Tools input files were processed using the PathoLogic module to produce population- and community-level ePGDBs. Average resource usage for pathway inference was 228.45 ± 110.04 mins / 1.92 ± 0.10 GBs, with the maxima and minima at 470.40 mins / 2.05 GBs and 112.70 mins / 1.77 GBs, respectively.

### 3.2 Usage, Accessibility, and Availability

Two significant barriers to adoption of prior versions of MetaPathways were the configuration file and installation procedure. A configuration file necessitates an understanding of all parameters before usage, including those for optional features that are not relevant to every user. We have integrated the configuration file into the CLI with default values assigned to optional parameters, such that a user only needs to learn the features specific to their use case. Similarly, manual installation procedures for reference databases impose a separate learning curve for the underlying local alignment tool (*e*.*g*., BLAST/FAST) in addition to that of MetaPathways. To mitigate the difficulty of installing reference databases, we developed a Snakemake ([27]) workflow, automating the installation of a select list of reference databases. The workflow also serves as an example for how to add additional reference databases should a user want to design and build a custom database. These changes lowered the learning curve “activation barrier” for new users compared to previous versions, making MetaPathways v3.5 more accessible.

MetaPathways v3.5 is open source software, and can be installed *via* the package manager Anaconda or as a container *via* Docker or Apptainer (formerly Singularity). Anaconda provides a standard channel for distributing data science tools and reduces the technical burden on users by automating the installation process. Some users may prefer a self-contained package for cloud and high-performance computing environments, hence both Docker and Apptainer containers are available for MetaPathways v3.5 at https://quay.io/repository/hallamlab/metapathways. We do not bundle the reference databases with the software due to licensing restrictions and size limitations, but it is automated to minimize the number of steps new users need to execute. The source code is available at https://bitbucket.org/BCB2/metapathways.

## 4 Conclusion

MetaPathways v3.5 supports scalable analysis of environmental genomes in an automated, interoperable, and now platform-agnostic workflow. New programmatic features improve Meta Pathways v3.5 accessibility for new users by abstracting away the technical complexities of both the installation and operation. As future expansions are planned, the underlying module-based infrastructure will continue to provide a flexible organizing principle to tame the growing complexity of Meta Pathways. The continued encapsulation of workflow complexity to maintain the accessibility achieved in this release will require more intelligent orchestration mechanisms for compute modules. We plan to add this feature by making future versions of Meta-Pathways more aware of interdependencies between modules. This will ensure that users can spend more time focused on interpreting MetaPath-ways outputs and less time setting up or managing large-scale workflows.

## Acknowledgements

**Funding:** This work was carried out under the auspices of the Digital Research Alliance of Canada formerly known as Compute/Calcul Canada, Genome Canada, Genome British Columbia, Genome Alberta, the Natural Science and Engineering Research Council (NSERC) of Canada, and the Canadian Foundation for Innovation (CFI) through grants awarded to SJH. ASH was supported in part by the Alexander Graham Bell Canada Graduate Scholarships-Doctoral Program (CGS D) and ECOSCOPE postdoctoral award administered by NSERC. CML was supported in part by a Mitacs Accelerate Fellowship postdoctoral award.

